# miRNova: A Next-Generation Platform for Ultra-Precise and Highly Specific MicroRNA Quantification Integrating a Tailored Stem-Loop RT-qPCR and a Robust Analytical Framework

**DOI:** 10.64898/2026.04.01.715903

**Authors:** Marc Van Der Hofstadt, Juliette Houot-Cernettig, Carolyn Thibal, Hoang Son Nguyen, Clément Marcelin, Alimata Ouedraogo, Pierre Champigneux, Laurence Molina, Malik Kahli, Franck Molina, Thi Nhu Ngoc Van

## Abstract

MicroRNAs (miRNAs) are ultra-short RNA molecules characterized by high sequence homology, frequent post-transcriptional modifications, and typically low abundance, particularly in circulating biofluids. These inherent biological features present substantial technical challenges for RT-qPCR– based quantification. Consequently, the development of miRNA RT-qPCR assays has required architectural adaptations at the reverse transcription (RT) stage to generate extended cDNA templates, thereby enabling effective downstream quantitative PCR amplification.

One widely adopted approach involves the enzymatic addition of a poly(A) tail to the 3′ end of miRNAs, followed by poly(T)-primed universal reverse transcription, which has gained broad acceptance due to its perceived sensitivity and simplified workflow. However, independent experimental evidence indicates that this architecture does not consistently provide the level of specificity required for reliable single-nucleotide (SN) discrimination, particularly when quantifying low-abundance circulating miRNA targets, as demonstrated in our previous study. An alternative strategy relies on miRNA-specific reverse transcription using stem-loop priming has been equally well accepted. When generically generated, this approach offers certain improved specificity, but its performance in resolving single-nucleotide differences remains limited.

In this article, we employed precision engineering to maximize specificity for both reverse transcription and qPCR steps. By tailoring both primer design and reaction architecture to the specific sequence features of each miRNA, we enable robust single nucleotide discrimination among these ultra-short targets. Prototype of ten different miRNova assays quantifying miRNAs whose sequences are differed in various configurations were tested on synthetic miRNA targets. For miRNova assay validation, saliva samples were elite rugby players submitted to small RNA extraction, then RT-qPCR. Spike-in of synthetic targets was applied for each quantification point to characterized the sensitivity, specificity and accuracy of the assays. Comparative analysis was performed between miRNova and two commercially available kits on the same sample set. The obtained results show a superior performance of miRNova assays allowing for sensitive and accurate quantification of miRNAs in saliva samples. Altogether, this results in modular, reproducible assays optimized for low-abundance miRNA detection in challenging biofluids, including saliva, positioning the platform beyond existing sensitivity-focused solutions toward true diagnostic-grade specificity.

## Introduction

MicroRNAs (miRNAs) have emerged as a central class of regulatory molecules with significant promise as non-invasive biomarkers. Their stability in extracellular environments and their direct involvement in gene regulation make them particularly attractive for applications ranging from disease diagnosis to longitudinal monitoring (Nemeth et al., 2023). Among accessible biofluids, saliva has gained increasing attention as a diagnostic matrix due to its non-invasive collection, rapid turnover, and compatibility with repeated sampling in real-world settings. These features make saliva especially relevant for applications requiring dynamic monitoring, such as sports medicine, neurology, and chronic disease management (Nonaka and Wong, 2022).

Despite this strong biological and clinical potential, the translation of miRNA biomarkers into routine diagnostic use remains limited, largely due to technical challenges associated with their quantification. miRNAs are extremely short (~22 nucleotides), often differ by only a single nucleotide within families, and are typically present at low abundance in biofluids. These characteristics place stringent demands on analytical methods, particularly in terms of specificity and sensitivity (Saliminejad et al., 2019) (Van Der Hofstadt et al., 2024).

Reverse transcription quantitative PCR (RT–qPCR) is widely regarded as the reference method for nucleic acid quantification. However, its application to miRNAs requires specialized adaptations. Two main strategies have been developed: poly(A)-tailing followed by universal reverse transcription (Shi et al., 2012), and miRNA-specific stem–loop priming (Chen, 2005). While both approaches have been enabled sensitive detection, they remain fundamentally limited in their ability to reliably discriminate between closely related miRNA sequences, especially when single-nucleotide differences are involved (Van Der Hofstadt et al., 2024). In practice, this limitation can lead to signal cross-reactivity, reduced quantitative accuracy, and ultimately, compromised biomarker interpretation (Faraldi et al., 2019).

A key issue underlying these limitations is that specificity is often enforced at only one stage of the workflow, typically during PCR amplification. As a result, mismatched targets may still be reverse-transcribed and subsequently amplified, particularly in low-abundance contexts where stochastic effects and kinetic competition become significant. This problem is further exacerbated in complex biological matrices such as saliva, where background nucleic acids, inhibitors, and variable RNA content can amplify non-specific signals and reduce assay robustness (Van Der Hofstadt et al., 2024).

Compared to poly(A)-tailling method, stem-loop method proposes a double specific priming strategy, which has been extended to several variants to overcome the limited specificity of miRNA quantification (Honda and Kirino, 2015a) (Ding et al., 2019) (Yang et al., 2014) (Honda and Kirino, 2015b) (Androvic et al., 2017). However, these adaptations have not yet been validated in clinical samples.

In this context, there is a clear need for next-generation RT–qPCR strategies that integrate specificity across the entire workflow, from reverse transcription to amplification, while maintaining high sensitivity and efficiency. Addressing this need is essential not only for improving analytical performance but also for enabling the reliable detection of low-abundance, clinically relevant miRNAs in real-world samples. In this study, we introduce miRNova, a modular RT–qPCR platform designed to overcome these limitations through precision, sequence-informed assay engineering. Unlike conventional approaches, miRNova combines tailored stem–loop reverse transcription primers, oligo blockers and stabilizers associated with strategic incorporation of locked nucleic acids (LNAs); and optimized amplification conditions to enforce discrimination at multiple levels. This integrated design aims to enhance early-stage selectivity, limit off-target amplification, and preserve efficient signal generation. The performance of miRNova was evaluated across a range of increasingly complex conditions, from synthetic miRNA targets to saliva-derived RNA. Particular emphasis was placed on single-nucleotide discrimination, using challenging miRNA pairs that differ either in central or terminal positions. Comparative analyses against widely used commercial platforms were performed to demonstrate that miRNova achieves substantially improved specificity without compromising amplification efficiency, resulting in greater separation between matched and mismatched targets. Importantly, this performance advantage is maintained in saliva, where miRNova enables the detection of low-abundance miRNAs that remain undetectable using existing methods.

## Materials and Methods

### Study approval and saliva collection

30 players from French National Under-20 men’s rugby team and healthy participants were recruited according to the personal protection committee (CPP) with registered number 2024-A02331-46. And 23.00930.000169 (NCT06149351 on www.clinicaltrials.gov), respectively. All subjects were informed, signed and consented in accordance with the CPP prior to the recruitment. Saliva collection and analyses were performed with approved protocols. A total of two collections (one collection pre-training and one post-training) were performed on the same day. Approximately 2 mL of unstimulated saliva was collected from each participant and stored at 4°C, transported to the laboratory Sys2diag, Montpellier, France. All saliva samples were aliquoted, treated with Qiazol (Qiagen) and stored at −80°C until used.

### Synthetic miRNA Targets

Ten miRNAs, including has-Let7a-5p, has-Let7f-5p, has-Let7d-5p, has-miR-16-5p, has-miR-21-5p, has-miR-103a-3p, has-miR-107, has-miR-134a-5p, has-miR-148a-3p, and has- and miR-155-5p were selected according to their sequence configurations and clinical relevance. All the ten synthetic miRNAs were purchased from Integrated DNA technologies (IDT, Europe).

### miRNA extraction

Total small RNA was extracted from 250 µL of whole saliva using miRNeasy Serum/Plasma Kit (Qiagen) according to the manufacture instructions (except for the elution step, where the extracted salivary small RNA was collected in 20 μL nuclease free water). In brief, 250 µL of whole saliva from each participant treated with 1 mL of Qiazol solution stored at −80°C was thawed on ice, homogenized by vortexing. Chloroform purification was followed by RNA precipitation by isopropanol, which was then loaded into a miRNeasy column, where only small RNA fragments (<200 bp fragments) are retained following multiple washes. All samples were handled and processed by the same manner with an equally respected delay between sampling and extraction time. All extracted small RNA samples were quantified by Nanodrop One (Thermo Scientific, Wilmington USA), normalized to 50ng/µL and subjected to reverse transcription.

### miRNA Quantification by miRNova assays

#### Primer Design

For reverse transcription (RT) primers, miRNA-specific stem-loop primers were designed following the strategy described by Chen et al. (Citation). Briefly, each RT primer consisted of a universal stem-loop structure coupled to a six or eight-nucleotide sequence at the 3′ end, complementary to the last nucleotides at the 3’ end of the target miRNA. miRNA sequences were retrieved from miRBase, and primer specificity, melting temperature, and secondary structures were evaluated using OligoAnalyzer to minimize primer-dimer formation and cross-reactivity among homologous miRNAs. For miRNA pairs whose sequences are differed only by nucleotides situated at their 3’ end, a Lock Nucleic Acid (LNA) was introduced at the differences. A cocktail of small oligos (6-8 nucleotides) was designed to complement parts of miRNA targets to act as stabilizers for RT reaction.

qPCR forward primers were designed to complement specifically the targeted miRNAs without overlapping the part that is already used in the stem-loop RT primer. For miRNA pairs whose sequences are differed by only one or several nucleotides, LNAs were introduced at the differences. Blocker oligos consist of a conjugated 3’ Phosphate was also designed for assays that target miRNAs whose sequences are closely similar to others OFF target miRNAs. Different versions of universal reverse primers complementing the Loop part of the RT primer were tested. Tm of all oligos was equilibrated by the addition extra nucleotide at their 5’ end or the introduction of one or several LNAs. Sequences of all designed oligos are presented in table S2.

#### Reverse transcription

Reverse transcription was performed using reverse transcriptase from Takara - PrimeScript RT Reagent (RR037A). In brief, 10 µL RT reactions containing 6 µL of 1–1013 copies of synthetic miRNAs or 50-300 ng of extracted small RNA, 2 µL of 5X RT buffer (containing dNTPs and RNase inhibitor), reverse transcriptase (from 0.125 – 0.5 µL), and stem-loop RT primers with concentrations ranges from 50 nM to 2 µM. H2O was added to make up the 10 µL reaction when needed. For all endogenous miRNA quantification, synthetic miRNAs were added as positive controls and plate calibrators. All experiments were performed in duplicate. No-template and no-RT enzyme controls were systematically included to monitor contamination and/or genomic DNA carryover.

Multiple thermal cycling conditions were tested: Program 1 consists of 42°C for 60 min 42°C for 60 min, followed by enzyme inactivation at 85 °C for 5 min; Program 2 consists of 16 °C for 30 min, 50°C for 30 min, followed by enzyme inactivation at 85 °C for 5 min; Program 3 consists of 16°C-30min, followed by 60 cycles of (30°C for 30s, 42°C for 30s and 55°C for 1min) and terminated with an enzyme inactivation at 95°C-5min. Program 4 consists of an incubation overnight at 4°C followed by full Program 3 thermocycling.

#### qPCR amplification

qPCR reactions were performed in a final volume of 10 µL containing 2.8 µL of 40X diluted cDNA template, 1µL containing 1.5 µM of Forward primers, and from 0.135-07 µM of Reverse primer, 5 µL of 2X Luna Universal qPCR Master Mix (M3003E) or LightCycler® 480 SYBR Green I Master (cat. nos. 04887352001), H2O was added to make up the 10 µL reaction when needed. All experiments were performed in duplicate. No-template controls were systematically included.

Amplification thermocycling conditions consists of an initial denaturation step at 95 °C for 10 min, followed by 40 cycles of denaturation at 95 °C for 10 s and annealing/extension step. Different temperatures were tested for the annealing/extension step, ranging from 50 to 70°C with extension time ranging from 20 to 60 s.

### miRNA Quantification by miRCURY*®* assays

#### Reverse Transcription of miRNAs

miRCURY LNA RT Kit employing poly (A) polymerase for tailing RNA prior to an universal reverse transcription (RT) using poly T primer was purchased from Qiagen. 10 µL RT reactions were prepared in 96 well plates containing 2 µL of 5X reaction buffer, 1 µL 10X reverse transcriptase and 1 µL of synthetic or extracted small RNA at desired concentrations. The reactions were incubated in a peqSTAR 96X thermocycler (Ozyme, Montigny-le-Bretonneux France) at 42°C for 60 min, followed by a denaturation step at 85°C for 5 min and stored at −20°C until used. No template and no reverse transcriptase enzyme negative controls were included in each run. All samples and control conditions were run in duplicate.

For the analysis using 50 ng of extracted small RNA as input, all extracted samples were normalized to 50 ng/µL and then 1 µL was added to the RT-qPCR reaction, except for P5 and P6 samples whose concentration were inferior to 50 ng/µL, and hence higher volumes were required.

#### Quantitative PCR

miRCURY LNA SYBR Green PCR Kit and target specific primers (miRCURY LNA miRNA PCR assays) were purchased from Qiagen. 10 μl qPCR reactions were prepared in 384 multi-well plates containing 5 µL of qPCR master mix; 1 µL of corresponding primers (miRCURY LNA miRNA PCR Assay) and 3 µL of 10X diluted RT product. QPCR negative controls (No-template reactions) were included for each assay. All samples and control conditions were run in duplicate. Real-time qPCR thermal cycling reactions were performed on a LightCycler 480 (Roche, Meylan France) directed by the LightCycler 480 Software (version 1.5.1.62). Thermal cycling conditions were: Pre-incubation for 2 minutes at 95°C, 40 cycles of amplification (95°C for 10s, 56°C for 60s). Ct values were analysed by Abs Quant/2nd Derivative Max of the same software. No template negative controls were included in each run. All samples and control conditions were run in duplicate.

### miRNA Quantification by TaqMan® assays

#### Reverse Transcription of miRNAs

Reverse transcription (RT) was performed using the TaqMan® MicroRNA Reverse Transcription Kit (ref: 4366596, Thermo Fisher Scientific), following the manufacturer’s protocol. Briefly, 50 ng of total small RNA was used per reaction. RT reactions (15 µL final volume) contained 1× RT buffer, 0.25 mM each dNTP, 3.33 U/µL MultiScribe™ Reverse Transcriptase, 0.25 U/µL RNase inhibitor, and miRNA-specific stem-loop RT primers (TaqMan® MicroRNA Assays). For all endogenous miRNA quantification, synthetic miRNAs were added as positive controls and plate calibrators.

Thermal cycling was carried out on a peqSTAR 96X thermocycler (Ozyme, Montigny-le-Bretonneux France) under the following conditions: 30 min at 16 °C, 30 min at 42 °C, followed by enzyme inactivation at 85 °C for 5 min. cDNA samples were then held at 4 °C and stored at −20 °C until quantitative PCR analysis. No template and no reverse transcriptase enzyme negative controls were included in each run. All samples and control conditions were run in duplicate.

#### Quantitative PCR (qPCR)

Quantitative PCR was performed using the TaqMan® Universal Master Mix II, no UNG (ref: 4440040, Thermo Fisher Scientific) on a LightCycler 480 (Roche, Meylan France) directed by the LightCycler 480 Software (version 1.5.1.62). Each 20 µL PCR reaction contained 1.33 µL of RT product, 1× TaqMan® Universal Master Mix II (no UNG), 1× TaqMan® MicroRNA Assay (specific probe and primers), and nuclease-free water. Amplification was carried out under the following cycling conditions: initial denaturation at 95 °C for 10 min, followed by 40 cycles of 95 °C for 15 s and 60 °C for 60 s. No template negative controls were included in each run. All samples and control conditions were run in duplicate.

#### qPCR Data Analysis

Cycle threshold (Ct) values were automatically determined using automatic baseline and threshold settings. Negative controls showed no amplification or late Cq values (≥35), confirming absence of contamination. All replicates with a standard deviation greater than 0.3 Ct were excluded from analysis. All assays were performed on its target (ON target) and the one whose sequence shares the highest similarity with the ON target (OFF target). Specificity of all assays was calculated as ΔCt between the two detections: the ON and OFF target. The Limit of Detection (LOD) was calculated for all assay using their standard curves.

For endogenous miRNA quantification, relative expression levels were calculated using the 2^ΔCt or 2^ΔΔCt method. Normalization was performed using validated endogenous controls and/or the average Ct value of all detected signals.

#### Statistical Analysis

Scipy version 1.11.2 was used on python 3.10.4 to do statistical analysis. All statistical test were done using Mann-Whitney U (scipy.stats.mannwhitneyu) with asymptotic method (i.e. p-value calculated by comparing to normal distribution and hence correcting for ties). All statistical calculations show the mean values ± standard error of the mean.

## Results

### miRNova Enables Robust Single-Nucleotide Discrimination (SND)

Specificity of miRNA RT-qPCR is often defined by discrimination phase of early cycles, kinetic competition and polymerase fidelity for perfectly matched sequences whilst delay extension of mismatched ones. miRNova assays for SND was engineered based on two distinct sequence configuration models: (i) To discriminates between two miRNAs with with one nucleotide difference situated in the middle of the molecule such as Let7a and Let7f; (ii) To discriminates between two miRNAs with one nucleotide difference situated at the 3’ end of the molecule such as miR-103 and miR-107. Different priming strategies were employed for these two models. For each assay, amplification was performed using both the perfectly matched target (ON target) and the corresponding SN-mismatched sequence (OFF target). ΔCt between the ON and the SN-OFF target is calculated as a measure for assay specificity. Corresponding Qiagen assays employing poly(A)-tailling approach were used for comparative analysis.

For Let7a and Let7f configuration, the two assays share the same stem loop primer as their 3’ end share the same sequence. Therefore, discrimination between these two molecules can only be envisaged at the qPCR step. For this model, an LNA was employed on the forward primer at the SN position of the two assays. Results on Figure 1A shows that, the initial generic stem-loop design (miRNova-1) for Let7a displayed a lightly better specificity compared to Qiagen assay with ΔCt of 4.31 and 3.66, respectively. The employment of an LNA in the forward primer (miRNova-2) further increase the ΔCt to 7.5. Additional increase in specificity for Let7a assay (miRNova-3, ΔCt = 9.09) was achieved by the use of a blocker oligo which is fully complement to the OFF target Let7f while being conjugated with a phosphate at the 3’ end, no amplification could occur. miRNova-4 ΔCt gain (11.19) was acquired by primer concentration adjustment and the final optimized miRNova with a ΔCt of 13.41 was obtained from RT and qPCR thermocycling optimization. Altogether, this optimization strategy generates a significant increase in specificity of the Let7a assay, displaying a ΔCt of 13.41 compared 3.66 of the Qiagen one.

**Figure 1.**
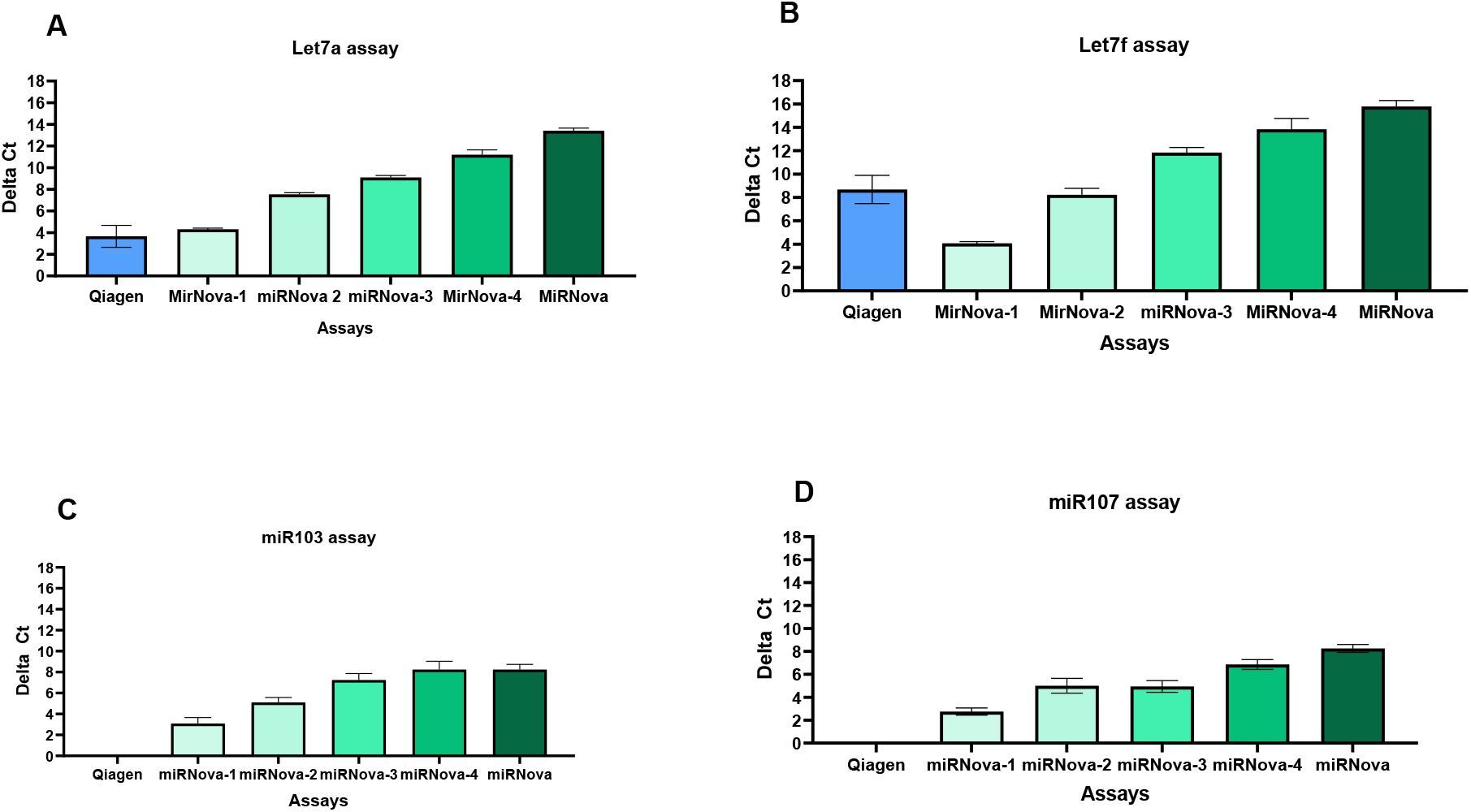
Specificity of optimized miRNova assays compared with Qiagen commercial kits using synthetic miRNA targets. Throughout the optimization process, miRNova assays increased their specificity resulting in greater differences in signal detection between ON and OFF target. **Panel A:** miRNova Let7a assay exhibits 3 times more specific (ΔCt =13.41) compared to Qiagen (ΔCt =3.66) kit. **Panel B:** miRNova Let7f assay is 2 times more specific (ΔCt =15.79) compared to Qiagen (ΔCt =8.64) kit. **Panel C:** miRNova miR-103 assay is capable of discriminating miR-107 target with ΔCt =8.23 while Qiagen kit detected the two targets at the same extend (ΔCt =0). **Panel D:** miRNova miR-107 assay is capable of discriminating miR-103 target with ΔCt =8.26 while Qiagen kit detected the two targets at the same extend (ΔCt =0). Specificity is represented as the mean ΔCt (Ct_OFF target − Ct_ON target) across all detectable concentrations, excluding data points within the saturation range (Ct ≥ 5). Standard Deviation was calculated from 3 independent experiments. Qiagen: miRCURY® assays, miRNova-1: Initial design prior to optimization. miRNova-2 – miRNova-4: Improving versions of miRNova. miRNova: Final optimized version. A high concentration (10^13^ copies) of synthetic target miRNAs was used for all presented experiments.

Results of Let7f assay in Figure 1B shows that, the initial design miRNova-1 displayed a poorer specificity compared with the Qiagen one, which was subsequently rescued by the introduction of an LNA to the SN position (miRNova-2). In the same line with Let7a assay, additional increases in specificity of the Let7f assay were achieved by employing blocker oligo (miRNova-3), primer concentration and RT/qPCR thermocycling optimization (miRNova-4), generating the final optimized version of miRNova assay for Let7f with a superior specificity reflecting by a ΔCt of 15.79 compared to 8.68 of the Qiagen one.

Contrary to Let7a and Let7f configuration, miR-103 and miR107 are differed by a SN situated at the 3’ end of the molecule, therefore their discrimination can only be achieved at the RT step. To this end, we employed an LNA in the SN position of the stem-loop primer accompanied with a stabilizer cocktail oligos. As shown in Figure 1C and 1D, the initial designs without LNA achieved a limited gain in specificity displaying a ΔCt of 3.08 and 2.75 for miR-103 and miR-107, respectively. However, these designs still overperformed their corresponding Qiagen assays whose detection signals were identical for both ON and OFF target, displaying a ΔCt of zero for both assays. The employment of a LNA in the stem-loop primer (miRNova-2) double the gain in ΔCt, which is additionally alimented by the stabilizer cocktails (miRNova-3), primer concentration (miRNova-4) and Thermo cycling optimization. The final optimized miRNova assay for both miR-103 and miR-107 achieves an a ΔCt of 8.23 and 8.28, respectively.

### miRNova Outperforms Commercial Platforms in Specificity Without Compromising Amplification Efficiency

The analytical performance of the optimized miRNova assays was then evaluated across a broad dynamic range of synthetic miRNA inputs, spanning from 1 to 10^13^ copies. This wide concentration gradient was selected to rigorously assess both sensitivity and linearity, as well as the ability of the assays to maintain specificity under conditions of extreme target abundance variation. To benchmark performance, miRNova assays were directly compared to widely used commercial platforms: the Qiagen system based on a poly(A)-tailing reverse transcription strategy, and the Thermo Fisher Scientific system combining stem–loop reverse transcription with TaqMan probe-based detection.

Two representative and challenging sequence configurations were selected for this comparative analysis: the Let-7a/Let-7f pair, which differs by a single nucleotide located in the central region of the sequence, and the miR-103/miR-107 pair, which differs at the 3′ end. These models allowed us to evaluate assay performance across distinct mismatch contexts that are known to differentially impact RT–qPCR specificity.

As illustrated in Figure 2, both commercial platforms exhibited intrinsic limitations. The Qiagen system, while robust in workflow simplicity, showed limited adaptability and consequently restricted analytical performance, particularly in terms of specificity (Fig. 2A, 2D, and 2G). In several cases, ON and OFF targets were poorly resolved, reflecting insufficient discrimination at the sequence level. The Thermo Fisher platform, which offers greater flexibility through probe-based detection, demonstrated modest improvements in specificity; however, this gain was accompanied by a noticeable reduction in amplification efficiency. This was evidenced by a deviation from ideal exponential amplification behavior, with amplification curves tending toward linearity at higher concentrations, suggesting suboptimal reaction kinetics.

**Figure 2.**
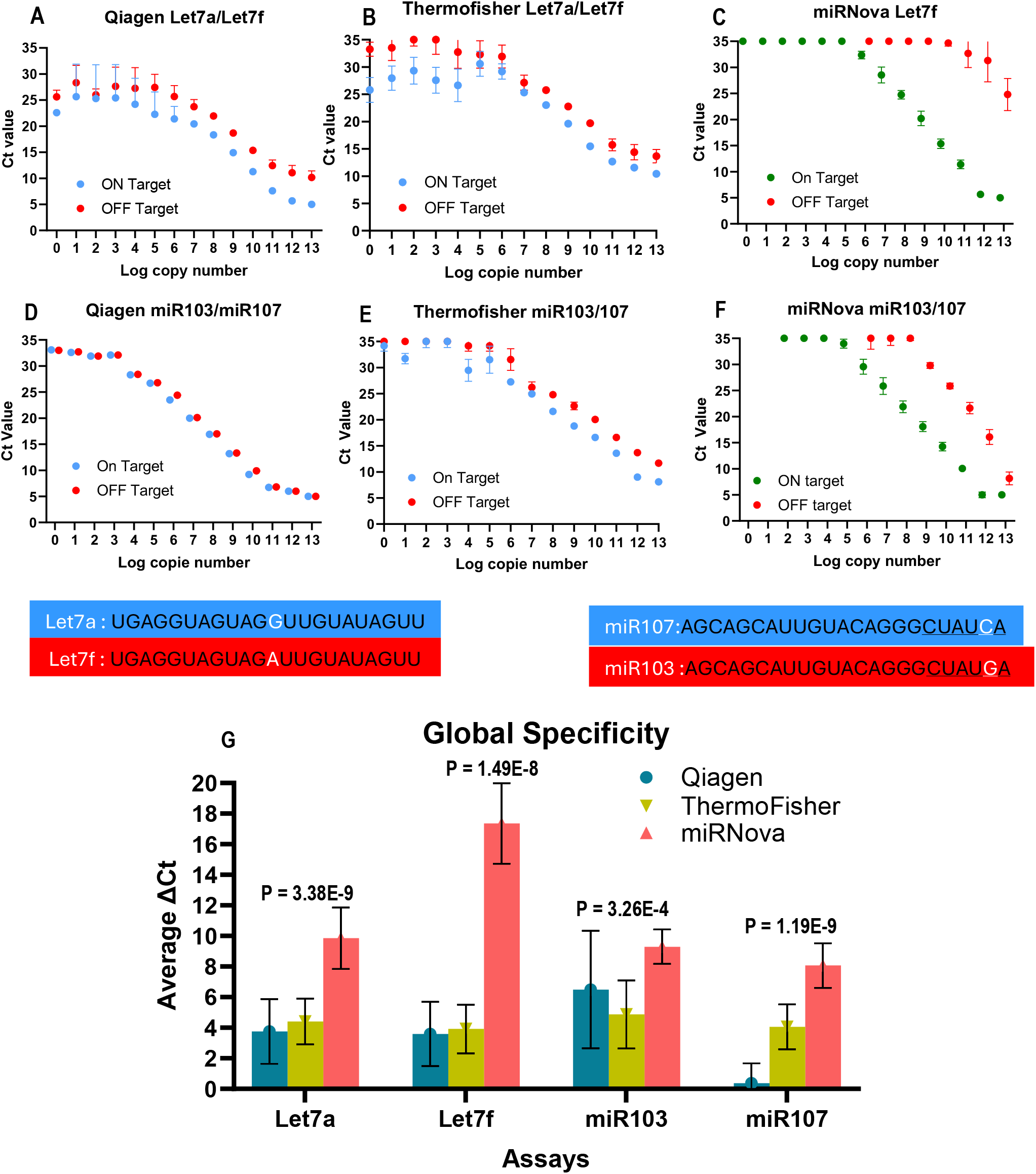
Comparative analytical performance of the miRNova platform versus Qiagen and Thermo Fisher Scientific systems using synthetic miRNA targets. The single-nucleotide (SN) discrimination capability of the miRNova platform was evaluated across two distinct sequence configurations: Let-7a versus Let-7f and miR-103 versus miR-107, in which the single-nucleotide difference is located in the central region and at the 3′ end of the molecules, respectively. **Panels A and D** show that Qiagen assays exhibit limited performance, characterized by low specificity and a high level of primer dimerization, resulting in relatively constant and elevated Ct values at low target concentrations. **Panels B and E** indicate that Thermo Fisher Scientific assays display a modest improvement in specificity; however, this is accompanied by a reduction in amplification efficiency, as evidenced by early signal saturation at Ct values higher than the theoretical minimum (Ct ≈ 9 rather than Ct ≈ 5). **Panel G** presents the global specificity metric, calculated as the mean ΔCt (Ct_OFF target − Ct_ON target) across all detectable concentrations, excluding data points within the saturation range (Ct ≥ 5). Statistical analysis was performed using Kruskal-Wallis method to compared the three platforms. Adjusted P values with Benjamini-Hochberg correction are presented.

In contrast, the miRNova platform was designed from the outset as a fully modular and tunable system, enabling precise adjustment of assay parameters according to the sequence characteristics of each target miRNA. This design strategy translated into consistently superior performance across all tested conditions. As shown in Figure 2C–E and summarized in Figure 2G, miRNova achieved both high amplification efficiency and excellent linearity over the entire concentration range, while simultaneously producing the largest Ct separation between perfectly matched (ON) targets and single-nucleotide mismatched (OFF) targets.

Importantly, this performance was not limited to a specific sequence configuration. The miRNova architecture proved robust across both central and 3′ mismatch models, highlighting its ability to accommodate diverse miRNA sequence constraints. This adaptability is a key advantage over existing fixed-design platforms, as it allows consistent optimization across heterogeneous miRNA targets.

Finally, the analytical gains observed with synthetic targets were subsequently confirmed in biological samples, including saliva—a highly complex matrix characterized by low RNA abundance and potential inhibitory factors. The ability of miRNova to maintain both sensitivity and specificity in such conditions underscores its robustness and real-world applicability, positioning it as a promising tool for high-fidelity miRNA quantification in clinical and translational settings.

These engineering models were then applied to develop other assays in various sequence configuration.

### miRNova Assay Optimization on biological samples

To further assess the practical applicability of the platform, we optimized miRNova assays for saliva, a particularly challenging biological matrix characterized by high biochemical complexity, variable composition, and extremely low miRNA abundance. This step was critical to evaluate assay performance under conditions where conventional RT–qPCR solutions often struggle to deliver reliable and reproducible results.

As shown in Figure 3A, both the Let-7a and Let-7f assays exhibited a clear dose-dependent response when increasing amounts of saliva-derived small RNA (50–300 ng) were used as input. The Let-7a assay demonstrated robust signal detection across this range, with Ct values decreasing from approximately 31 to 26, reflecting efficient amplification and good sensitivity. Similarly, the Let-7f assay showed consistent and reproducible detection, with Ct values ranging from ~33 to 29, remaining well within a reliable quantification window despite its slightly lower abundance or assay efficiency. Importantly, no evidence of reaction inhibition was observed across the tested RNA inputs for either assay, indicating that the miRNova chemistry is well tolerated in the presence of saliva-derived components that are often inhibitory in standard workflows.

**Figure 3.**
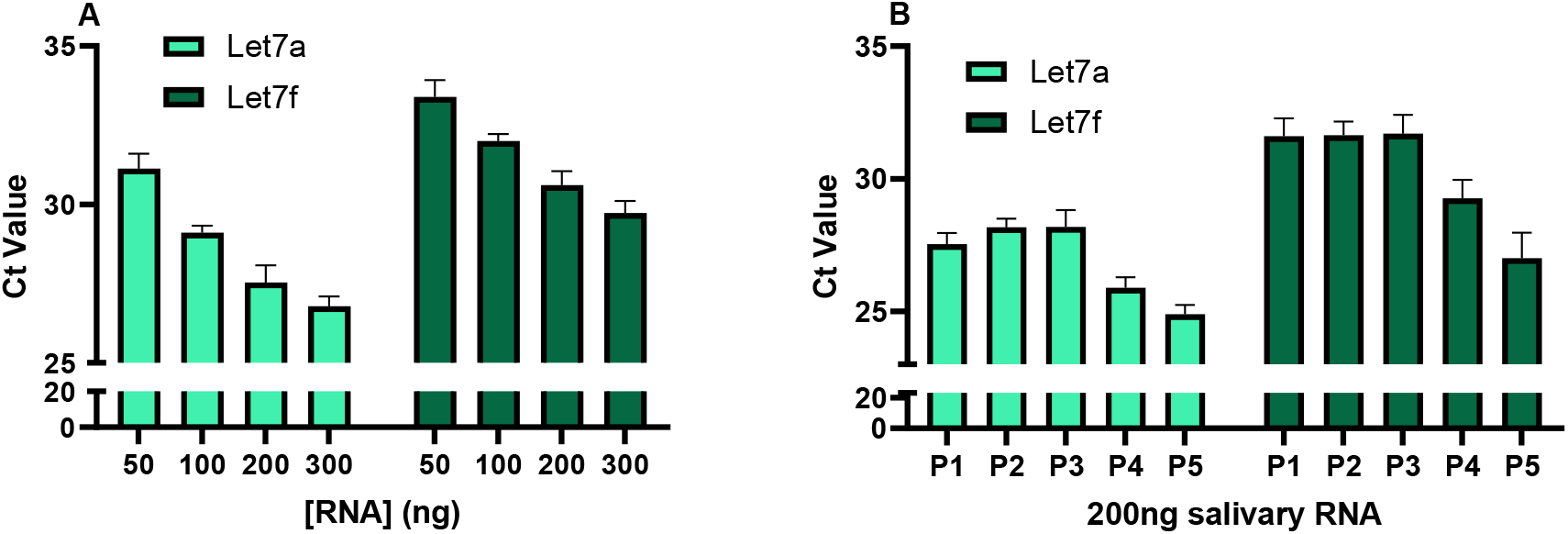
miRNova Let7a and Let7f assays detecting salivary miRNAs in saliva samples in optimized conditions. **Panel A:** Dose dependent signals of the two assays using 50-300 ng extract small RNA input. **Panel B:** Optimized assay efficiency by different RT thermos cycling programs. P1: 42°C for 60 min. P2: 16°C for 30 min followed by 42°C for 30 min. P3: 16°C for 30 min followed by 50°C for 30 min. P4: 16°C-30min followed by 60 cycle of: 30°C-30s; 42°C-30s; 55°C-1min. A pool of small RNA extracted from different saliva samples was normalized and used in all tested assay.

To further enhance detection sensitivity, we systematically evaluated several reverse transcription (RT) thermal cycling programs using a fixed input of 200 ng of saliva-derived small RNA (Figure 3B). Compared to the initial RT condition (Program P1), alternative protocols P2 and P3 did not yield measurable improvements in signal detection for either assay, suggesting that simple adjustments in incubation temperature or duration are insufficient to overcome the intrinsic limitations of RT efficiency in this context.

**Figure 3.**
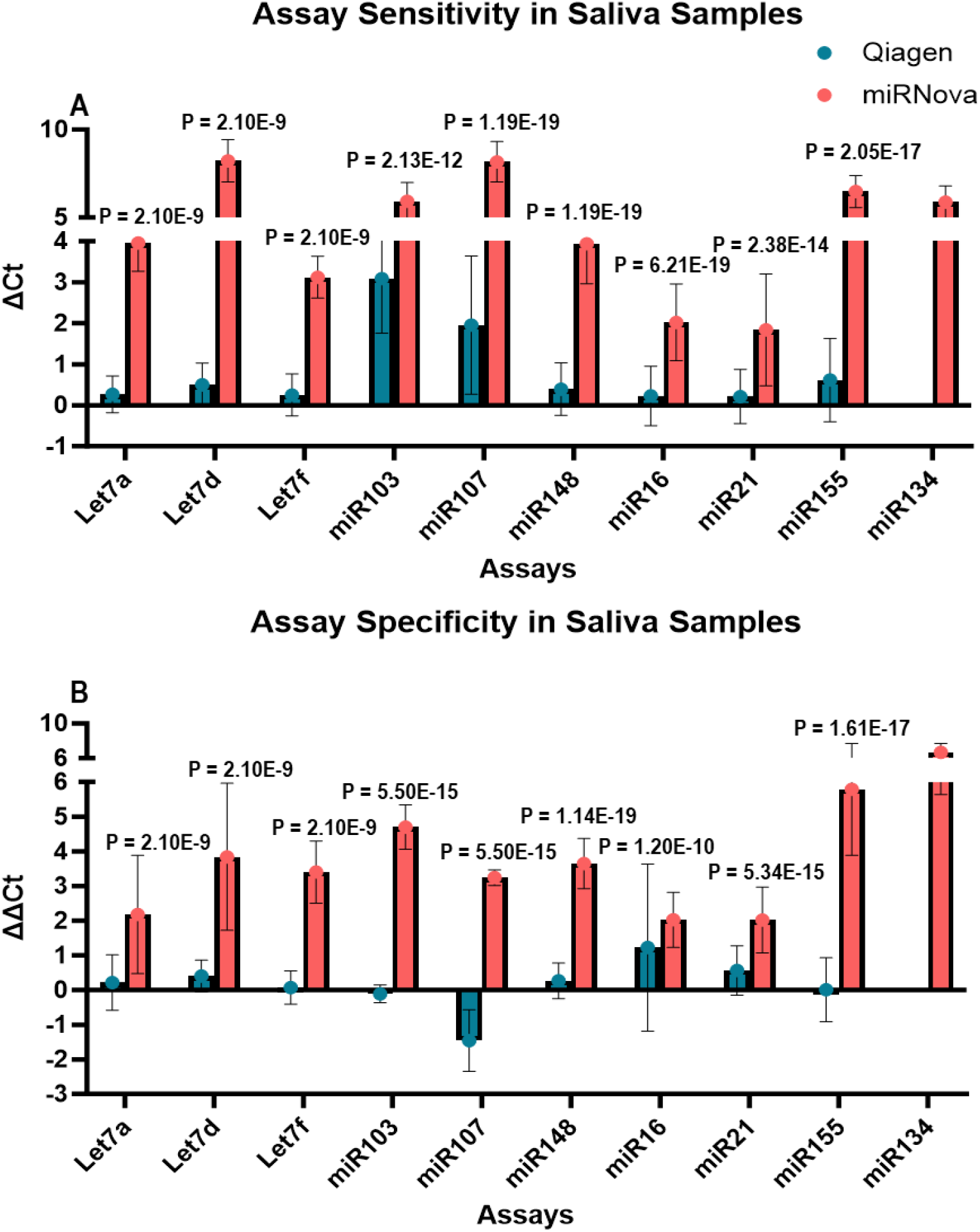
Comparative performance of the miRNova platform and the Qiagen system using endogenous small RNAs extracted from saliva samples. Small RNAs were isolated from 60 saliva samples obtained from 30 elite athletes and subsequently analyzed by RT-qPCR for the quantification of 10 miRNAs. To assess assay sensitivity and specificity in a biologically relevant context, 10^7^ copies of synthetic ON and OFF targets were spiked into the endogenous RNA samples. For each assay, sensitivity was defined as the response to the addition of the ON target (ON), whereas specificity was determined by correcting this response for cross-reactivity with the single-nucleotide-mismatched OFF target at the same concentration. **Panel A** shows assay sensitivity calculated as the ΔCt between the endogenous sample and the same sample spiked with the ON target (ΔCt_ON = Ct_Endo − Ct_Endo+ON). **Panel B** shows assay specificity expressed as ΔΔCt, defined as the difference between the sensitivity signal and the crosstalk generated by the OFF target (ΔCt_OFF = Ct_Endo − Ct_Endo+OFF; ΔΔCt = ΔCt_ON − ΔCt_OFF). Statistical analysis was performed using MannWithney method, adjusted P values are presented.

In contrast, more complex cycling strategies (Programs P4 and P5), which incorporate stepwise or iterative temperature transitions, resulted in substantial gains in assay performance. Notably, Program P5 provided the most significant improvement. Under these conditions, the Let-7a assay exhibited a Ct value of 24.89 compared to 27.54 with the initial program, corresponding to an approximate 3-cycle gain, indicative of a marked increase in effective cDNA yield. Even more pronounced improvements were observed for the Let-7f assay, where the Ct value decreased from 31.61 to 27.00, representing an approximate 4-cycle gain.

These improvements likely reflect enhanced primer–template hybridization dynamics and more efficient reverse transcription under cycling conditions, particularly for low-abundance targets embedded in complex RNA backgrounds. Overall, these results highlight the importance of RT step optimization as a key determinant of assay sensitivity, and demonstrate that the miRNova platform can be finely tuned to achieve robust and sensitive miRNA detection even in highly challenging biological matrices such as saliva.

### Integrated Performance Demonstrates Clinical-Grade Analytical Robustness

To further challenge the robustness of the platform under real-world conditions, we evaluated miRNova performance in saliva—arguably one of the most demanding biological matrices due to its intrinsic complexity, variable composition, and the extremely low abundance of circulating miRNAs. This experimental setting was specifically chosen to test the platform in conditions where conventional RT–qPCR approaches often reach their technical limits.

A panel of ten miRNova assays was applied to a cohort of sixty saliva samples, and their performance was assessed in parallel with corresponding assays from Qiagen under identical experimental conditions. To obtain a rigorous and biologically relevant evaluation of both sensitivity and specificity, each endogenous RNA sample was supplemented with 10^7^ copies of synthetic miRNA targets, including both perfectly matched sequences (ON targets) and single-nucleotide mismatched counterparts (OFF targets). Within this framework, assay sensitivity was defined as the magnitude of signal increase following the addition of the ON target, while specificity was assessed by correcting this response for any signal contribution arising from OFF-target amplification at the same concentration.

As shown in Figure 3, miRNova demonstrated consistently superior performance across all assays and samples. Notably, the platform successfully detected all ten miRNAs included in the panel, including miR-134, a low-abundance target that remained undetectable using the comparator system. This result is particularly significant, as it highlights the ability of miRNova to extend the detectable biomarker space in complex biological samples, an essential requirement for clinical applications.

Quantitative analysis of sensitivity further reinforced this advantage. Upon spike-in of ON targets, miRNova produced markedly stronger signal responses across all assays compared to the Qiagen platform (Figure 3A), indicating more efficient target capture and amplification. This enhanced sensitivity was observed consistently across the entire sample cohort, suggesting that the improvement is not sample-dependent but rather intrinsic to the assay design.

Importantly, this gain in sensitivity did not come at the expense of specificity. miRNova maintained robust discrimination between ON and OFF targets, even within the complex background of saliva-derived RNA. The ability to preserve high ΔΔCt values under these conditions demonstrates that the platform effectively limits cross-reactivity and maintains sequence-level resolution, a key requirement for accurate miRNA profiling.

Taken together, these results establish miRNova as a high-performance, saliva-compatible RT–qPCR platform, capable of delivering both sensitive and specific miRNA detection in challenging biological contexts. Its consistent performance across multiple targets and samples underscores its potential for reliable deployment in clinical and translational applications, particularly in non-invasive diagnostic settings.

## Discussion

Accurate quantification of microRNAs remains one of the most technically challenging tasks in molecular diagnostics. This difficulty arises from a unique combination of factors: the extremely short length of miRNAs, their high degree of sequence homology, and their often very low abundance in clinically relevant biofluids. While RT–qPCR has long been considered the gold standard for nucleic acid quantification, its direct application to miRNAs has required substantial methodological adaptations— many of which introduce trade-offs between sensitivity, specificity, and robustness.

In this study, we present miRNova, a platform designed to explicitly address these trade-offs through a precision-engineered, sequence-informed architecture. Rather than relying on generic or universal workflows, miRNova adopts a tailored strategy in which each assay is optimized according to the structural and thermodynamic properties of the target miRNA. This design philosophy proved essential to achieving one of the most critical—and historically difficult—objectives in miRNA analysis: reliable single-nucleotide discrimination.

### Revisiting the Specificity–Sensitivity Trade-Off

The major limitation of existing miRNA RT–qPCR methods lies in their inability to simultaneously maximize specificity and amplification efficiency. Poly(A)-tailing approaches, while convenient and often sensitive, rely on universal reverse transcription that inherently lacks sequence-level discrimination at the RT step. As a result, specificity must be enforced during qPCR alone, where the window for discrimination is narrower and more prone to kinetic leakage. This limitation becomes particularly evident when distinguishing closely related miRNAs, where even a single mismatch may not sufficiently delay amplification (Van Der Hofstadt et al., 2024).

Conversely, stem–loop-based methods introduce a degree of specificity during reverse transcription, yet their performance is highly dependent on primer design. Generic implementations often fail to fully exploit this potential, especially when mismatches occur at the 3′ end of the miRNA—an area known to be challenging due to its critical role in primer extension.

miRNova addresses these issues by synchronizing specificity across both RT and qPCR stages. By precisely controlling primer–target interactions—through adjustable complementarity length, strategic placement of locked nucleic acids (LNAs), and the use of auxiliary stabilizing and blocking oligonucleotides—the platform reinforces discrimination early in the reaction. This early-stage selectivity is key: mismatched targets are less likely to be reverse-transcribed efficiently, reducing their contribution to downstream amplification and thereby increasing ΔCt separation.

Importantly, this enhanced specificity is achieved without sacrificing amplification efficiency, a common drawback in highly stringent systems. The preservation of near-ideal amplification kinetics observed with miRNova suggests that the platform successfully balances thermodynamic stringency with enzymatic compatibility.

### Mechanistic Basis of Improved Single-Nucleotide Resolution

The ability of miRNova to discriminate single-nucleotide differences across both central and 3′ mismatch configurations provides important mechanistic insights. In conventional systems, discrimination efficiency is highly context-dependent, with central mismatches generally easier to resolve than terminal ones. The fact that miRNova performs robustly in both scenarios indicates that specificity is not solely governed by primer binding at a single site, but rather by a distributed network of interactions. Several elements likely contribute to this effect: (i) LNA incorporation increases local binding affinity and sharpens mismatch penalties, effectively amplifying the energetic difference between matched and mismatched hybrids; (ii) Shortened and tunable RT primer complementarity reduces tolerance to mismatches, particularly at critical positions; (iii) Stabilizer oligonucleotides may facilitate correct folding and hybridization dynamics during reverse transcription, improving efficiency for ON targets while maintaining discrimination; (iv) Blocker oligos actively suppress amplification of closely related OFF targets, introducing an additional layer of selectivity during qPCR.

Together, these features create a system in which specificity is enforced at multiple levels— thermodynamic, kinetic, and competitive—resulting in a cumulative amplification of discrimination that exceeds what can be achieved by any single mechanism alone.

### Performance in Complex Biological Matrices

One of the most compelling aspects of this study is the demonstration of miRNova’s performance in saliva, a notoriously challenging biological matrix (Lee and Wong, 2009) (Nonaka and Wong, 2022) (Ostheim et al., 2020). Unlike synthetic systems, saliva introduces a range of confounding factors, including enzymatic degradation, inhibitors, variable RNA content, and background nucleic acids. These factors often exacerbate non-specific amplification and reduce assay sensitivity, generating controversial and study dependent data (Hiskens et al., 2022).

Despite these challenges, miRNova maintained both high sensitivity and strong specificity, successfully detecting all targeted miRNAs across a cohort of 60 samples. Notably, the platform enabled detection of low-abundance miRNAs that were not measurable using comparator technologies. This finding is particularly significant, as the clinical utility of miRNA biomarkers often depends on the reliable detection of subtle expression changes in low-copy-number targets (Hindson, 2023) (Mavroudis et al., 2023) (Mavroudis et al., 2024) (Setti et al., 2020).

The robustness observed here suggests that miRNova is not only analytically superior under controlled conditions but also operationally resilient in real-world settings. This is a critical requirement for translation into clinical and point-of-care applications.

### Modularity as a Key Advantage for Translational Applications

Beyond performance metrics, a defining feature of miRNova is its modular design. Unlike fixed commercial platforms, which offer limited flexibility, miRNova allows each assay to be adapted based on the specific sequence characteristics of the target miRNA. This adaptability is particularly valuable given the diversity of miRNA families and the prevalence of closely related isoforms. From a translational perspective, this modularity offers several advantages: (i) Custom assay development for emerging biomarkers; (ii) Scalability across panels of miRNAs with heterogeneous sequence properties; (iii) Compatibility with different sample types and input conditions. Such flexibility is essential for bridging the gap between biomarker discovery and clinical deployment, where assay requirements often evolve rapidly.

### Limitations and Future Perspectives

While the results presented here are highly encouraging, several aspects warrant further investigation. First, although the platform was validated across a panel of ten miRNAs, broader validation across larger and more diverse miRNA repertoires will be important to confirm generalizability. Second, while saliva provides a stringent test environment, extending validation to other clinically relevant biofluids (e.g., plasma, cerebrospinal fluid) would further strengthen the platform’s applicability.

In addition, future work could explore automation and standardization of assay design, potentially integrating computational tools to streamline primer optimization. This would facilitate wider adoption and reduce the expertise barrier associated with highly tailored assay development.

Finally, integration with multiplexing strategies could further enhance throughput and quantitative precision, opening new avenues for clinical diagnostics and large-scale screening.

## Conclusion

miRNova represents a significant step forward in miRNA quantification technology. By addressing the fundamental limitations of existing RT–qPCR approaches through multi-layered, sequence-informed design, the platform achieves a rare combination of high specificity, strong sensitivity, and robustness in complex biological matrices.

These characteristics are essential for unlocking the full potential of miRNAs as clinical biomarkers. As such, miRNova not only advances the technical state of the art but also provides a practical pathway toward reliable, non-invasive molecular diagnostics.

## Acknowledgements

The authors would like to thank all participants who made this study possible. We sincerely thanks Victor Petit, CEO of the company SkillCell for all the insightful discussion and comments from the beginning till the manuscript writing process. We thanks the CNRS for being the promotor of the two clinical studies.

## Author contributions

Study concept and design: M.VDH & TNN.V. Acquisition, analysis, interpretation of data: All authors. Writing of the manuscript and critical revision for important intellectual content: M.VDH., F.M. and TNN.V. Statistical analysis: M.VDH&& TNN.V. Funding acquisition: F.M and M.VDH. All authors critically reviewed and edited the manuscript.

## Data availability statement

The authors declare that all relevant data has been provided within the manuscript and its supporting information files.

